# Highly sensitive and full-genome interrogation of SARS-CoV-2 using multiplexed PCR enrichment followed by next-generation sequencing

**DOI:** 10.1101/2020.03.12.988246

**Authors:** Chenyu Li, David N. Debruyne, Julia Spencer, Vidushi Kapoor, Lily Y. Liu, Bo Zhou, Utsav Pandey, Moiz Bootwalla, Dejerianne Ostrow, Dennis T Maglinte, David Ruble, Alex Ryutov, Lishuang Shen, Lucie Lee, Rounak Feigelman, Grayson Burdon, Jeffrey Liu, Alejandra Oliva, Adam Borcherding, Hongdong Tan, Alexander E. Urban, Xiaowu Gai, Jennifer Dien Bard, Guoying Liu, Zhitong Liu

**Affiliations:** Paragon Genomics Inc., Hayward, CA 94545 USA; Department of Psychiatry and Behavioral Sciences, Department of Genetics, Stanford University, CA 94305 USA; MGI, BGI-Shenzhen, Shenzhen 518083 China; BGI-Shenzhen, Shenzhen 518083 China; Department of Pathology and Laboratory Medicine, Children’s Hospital Los Angeles, Los Angeles, CA 90027

## Abstract

Many detection methods have been used or reported for the diagnosis and/or surveillance of COVID-19. Among them, reverse transcription polymerase chain reaction (RT-PCR) is the most commonly used because of its high sensitivity, typically claiming detection of about 5 copies of viruses. However, it has been reported that only 47-59% of the positive cases were identified by some RT-PCR methods, probably due to low viral load, timing of sampling, degradation of virus RNA in the sampling process, or possible mutations spanning the primer binding sites. Therefore, alternative and highly sensitive methods are imperative. With the goal of improving sensitivity and accommodating various application settings, we developed a multiplex-PCR-based method comprised of 343 pairs of specific primers, and demonstrated its efficiency to detect SARS-CoV-2 at low copy numbers. The assay produced clean characteristic target peaks of defined sizes, which allowed for direct identification of positives by electrophoresis. We further amplified the entire SARS-CoV-2 genome from 8 to half a million viral copies purified from 13 COVID-19 positive specimens, and detected mutations through next generation sequencing. Finally, we developed a multiplex-PCR-based metagenomic method in parallel, that required modest sequencing depth for uncovering SARS-CoV-2 mutational diversity and potentially novel or emerging isolates.

## Introduction

A variety of methods for detecting SARS-CoV-2 have been reported and discussed^1,2^, including RT-PCR, serological testing^3^ and reverse transcription-loop-mediated isothermal amplification^4,5^. Currently, RT-PCR is considered the gold standard for diagnosing SARS-CoV-2 infections because of its ease of use and high sensitivity. RT-PCR has been reported to detect SARS-CoV-2 in saliva^6^, pharyngeal swab, blood, rectal swab^7^, urine, stool^8^, and sputum^9^. In laboratory conditions, the RT-PCR methodology has been shown to be capable of detecting 4-8 copies of virus or lower, through amplification of targets in the Orf1ab, E and N viral genes, at 95% confidence intervals^10-12^. However, only about 47-59% of the true positive cases were identified by RT-PCR, and 75% of RT-PCR negative results were actually later found to be positive with other assays, hence mandating repeated testing ^8,13-15^. In addition, there is evidence suggesting that heat inactivation of clinical samples causes loss of viral particles, thereby hindering the efficiency of downstream diagnosis^16^.

Therefore, it is necessary to develop robust, sensitive, specific and highly quantitative methods for reliable diagnostics^17,18^. The urgency to develop an effective surveillance method that can be easily used in a variety of laboratory settings is underlined by the wide and rapid spreading of SARS-CoV-2^19-21^. In addition, such method should also distinguish SARS-CoV-2 from other respiratory pathogens such as influenza virus, parainfluenza virus, adenovirus, respiratory syncytial virus, rhinovirus, human metapneumovirus, SARS-CoV, etc., as well as *Mycoplasma pneumoniae, Chlamydia pneumoniae* and other causes of bacterial pneumonia^22-25^. Furthermore, obtaining full-length viral genome sequence through next generation sequencing (NGS) prove to be essential for the surveillance of SARS-CoV-2’s evolution and for the containment of community spread^26-29^. Indeed, SARS-CoV-2 phylogenetic studies through genome sequence analysis have provided better understanding of the transmission origin, time and routes, which has guided policy-making and management procedures^27,28,30-33^.

Here, we describe the development of a highly sensitive and robust detection assay incorporating the use of multiplex PCR technology to identify SARS-CoV-2 infections. Theoretically, the multiplex PCR strategy, by simultaneously targeting and amplifying hundreds of targets, has significantly higher sensitivity than RT-PCR and may even detect nucleotide fragments resulting from degraded viral genomes. Multiplex PCR has been shown to be an efficient and low-cost method to detect Hantaan orthohantavirus and *Plasmodium falciparum* infections^34,35^, with high coverage (median 99%), specificity (99.8%) and sensitivity. Moreover, this solution can be tailored to simultaneously address multiple questions of interest within various epidemiological settings^35^. Similar to a recently described metagenomic approach for SARS-CoV-2 identification^36^, we also establish a user-friendly multiplex-PCR-based metagenomic method that is capable of detecting SARS-CoV-2, and could also be applied for the identification of novel pathogens with a moderate sequencing depth of approximately 1 million reads.

## Results

### Mathematical model of RT-PCR

Several RT-PCR methods for detecting SARS-CoV-2 have been reported to date^6,10-12^. Among them, two groups reported the detection of 4-5 copies of the virus^10,12^. To investigate the opportunity for further improvement upon the sensitivity of RT-PCR, we built a mathematical model to estimate the limit of detection (LOD) for SARS-CoV-2. The reported RT-PCR amplicon lengths are around 78-158bp, and the SARS-CoV-2 genome is 29,903bp (NC_045512.2). Thus, we chose 100bp amplicon length and 30kb SARS-CoV-2 genome size for mathematical modeling. With the assumption of 99% RT-PCR efficiency^11^, we found that RT-PCR assays could only detect 4.8 copies of SARS-CoV-2 at 95% probability (Fig. 1A), which is consistent with the experimental results previously reported^10^. In this model, the predicted probability of RT-PCR assays to detect one copy of SARS-CoV-2 is only 26% (Supplemental Fig. 1). This finding may explain, at least in part, the reported 47-56% detection rates of SARS-CoV-2 with known positive samples by RT-PCR^8,13^. We further discovered that the LOD appears to be independent of the viral genome size. For genomes of 4 to 100kb, the detection limit remains 4.8 copies at 95% probability.

**Figure 1.**
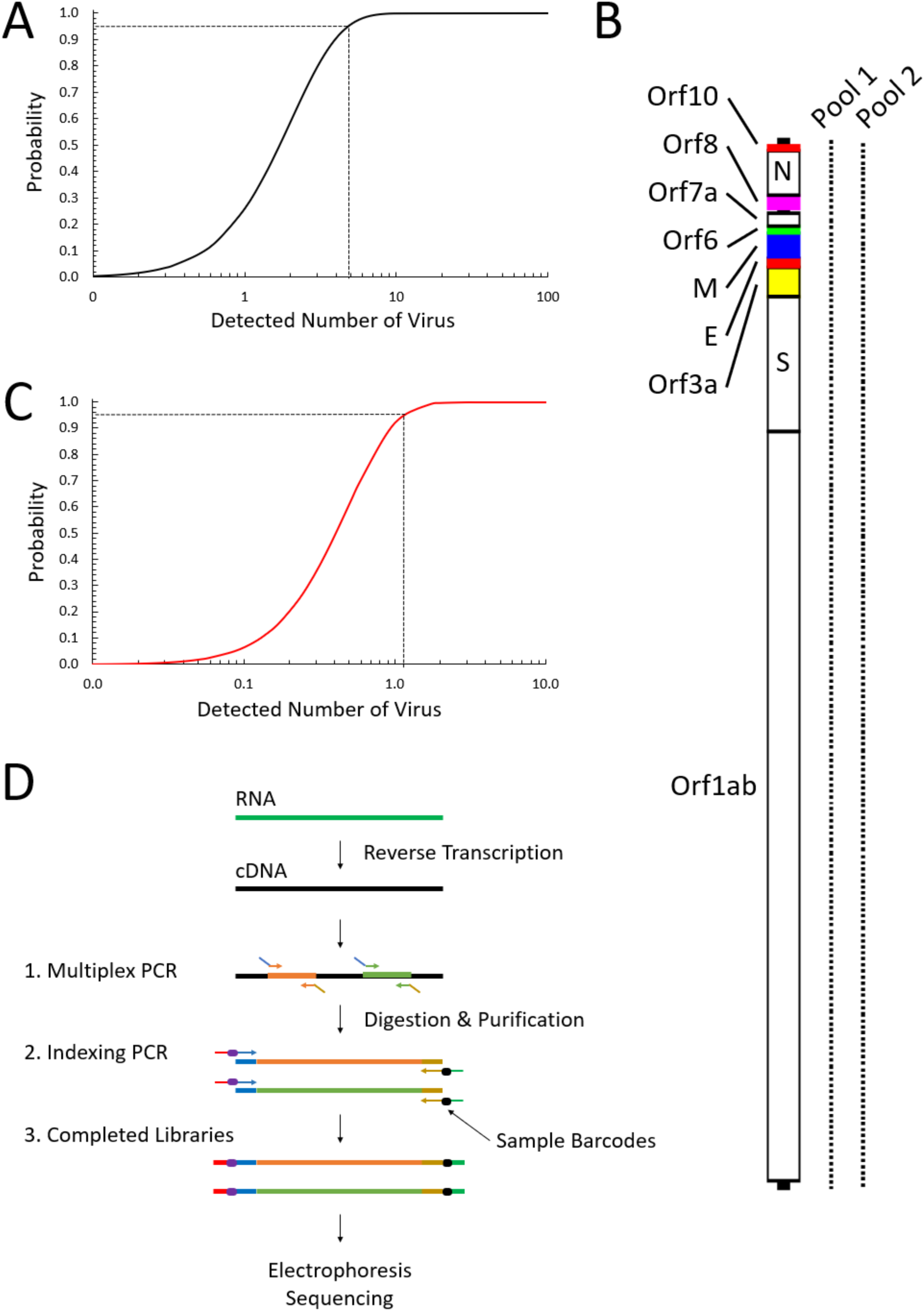
Mathematical model, primer design and workflow. (**A**) A mathematical model of RT-PCR based on Poisson process. The LOD is 4.8 copies of virus at 95% probability. (**B**) Two overlapping pools of multiplex PCR primers, shown on the right of the genome of SARS-CoV-2, were designed to span the entire viral genome. Pool 1, containing 172 pairs of primers, covers 56.9% of the viral genome and was used in the detection. Pool 2 contains 171 pairs of primers and covers 56.4% of the genome. Both pools are used to cover the full length of the genome. (**C**) A mathematical model of multiplex PCR with pool 1 of the primers. The LOD is 1.15 copies of virus at 95% probability. (**D**) The workflow of the multiplex PCR method. The prepared libraries can be detected using high-resolution electrophoresis, and sequenced together with other samples using high-throughput sequencing.

One way to elevate the sensitivity is to simultaneously target and amplify multiple regions of the viral genome in a multiplex PCR reaction, thereby increasing the frequency of occurrence in the mathematical model. Amplifying multiple targets has the advantage of potentially detecting fragments of degraded viral nucleotide fragments while tolerating genomic variations, thus allowing for the detection of new and ever-evolving viral strains. The amplification efficiency of multiplex PCR is critical for LOD. We estimated that the efficiency of our multiplex PCR technology is about 26% if using Unique Molecular Identifier (UMI)-labeled primers to count the amplified products after NGS sequencing (Supplemental Fig. 2 and Supplemental Table 1). However, the amplification efficiency could be lower, and amplicons would not be equally amplified if the template used is one single strand of cDNA. Thus, more amplicons are potentially required for multiplex PCR to detect limited copies of the virus.

### Mathematical model of multiplex-PCR-based detection method

We designed a panel of 172 pairs of multiplex PCR primers in order to increase the sensitivity of detecting SARS-CoV-2 (Fig. 1B). The average amplicon length is 99bp. The amplicons span across the entire SARS-CoV-2 genome with an average 76bp gap (76±10bp) between adjacent amplicons. Since the observed efficiency of multiplex PCR is about 26% in amplifying the four DNA strands of a pair of human chromosomes, we assumed an efficiency of 6% in amplifying the single-strand of a cDNA molecule. In addition, it has already been reported that 79% of variants are recovered when directly amplifying 600 amplicons from a single cell using our technology^37^. Therefore, we assumed that 80% of targeted regions would be amplified successfully. Using the same mathematical model described above, we estimated that our SARS-CoV-2 panel can detect 1.15 copies of the virus at 95% probability (Fig. 1C). Again, the LOD is independent of virus genome size.

We also designed a second pool of 171 multiplex PCR primer pairs. The target regions of these primer pairs overlap with the gaps between target regions of the previous pool of 172 primer pairs (Fig. 1B). Together, these two overlapping pools of primers provide full coverage of the entire viral genome. Most importantly, using both pools in detection would lead to a calculated detection limit of 0.29 copies at 95% probability.

### Detecting limited copies of SARS-CoV-2

The workflow was designed so that the multiplex PCR products are further amplified in a secondary PCR, during which sample indexes and NGS sequencing primers are added (Fig. 1D). The PCR products were first analyzed by electrophoresis to visualize potential positives. Since dozens of target regions could be amplified from a single copy of SARS-CoV-2, electrophoresis peaks with a defined distribution of peak sizes were expected. Multiplex PCR could potentially amplify not only SARS-CoV-2, but also other coronaviruses, due to shared sequence similarities despite the fact that we designed primers specific to the SARS-CoV-2 genome to avoid cross amplification. In that context, electrophoresis analysis provides a fast and sensitive indication of infection from at least that family of viruses. For specificity, the generated NGS sequencing library can be interrogated for definitive identification of the specific virus.

Two plasmids, containing the full sequence of the S and N genes of SARS-CoV-2, respectively, were used to validate our multiplex PCR method. A total of 28 targets are expected to be amplified within our 172-amplicon panel. To simulate the use of real clinical samples, these two plasmids were spiked into cDNA generated from human total RNA. The copy number of each plasmid was precisely determined by droplet-based digital PCR (ddPCR) with a QX200 system from Bio-Rad^38^. The two plasmids were diluted from approximately 9,000 copies to below one copy, and were amplified in multiplex PCR reactions. The library peaks of expected sizes were obtained from 8,900 to 2.8 copies of plasmids (Fig. 2A). Quantification of peaks demonstrated a wide dynamic range from 1 to about 1,000 copies of plasmids (Fig. 2B). The yield of the libraries started to saturate when the copy number reached 1,700. It is possible that the saturation point could be even lower when all of the 172 amplicons are amplified from positive clinical COVID-19 samples, and the library peaks could be observed with even fewer viral copies. In contrast, the detected quantities of a single target on S gene by RT-PCR rapidly dropped when using 2.85 copies (Fig. 2B and Supplemental Fig. 3).

**Figure 2.**
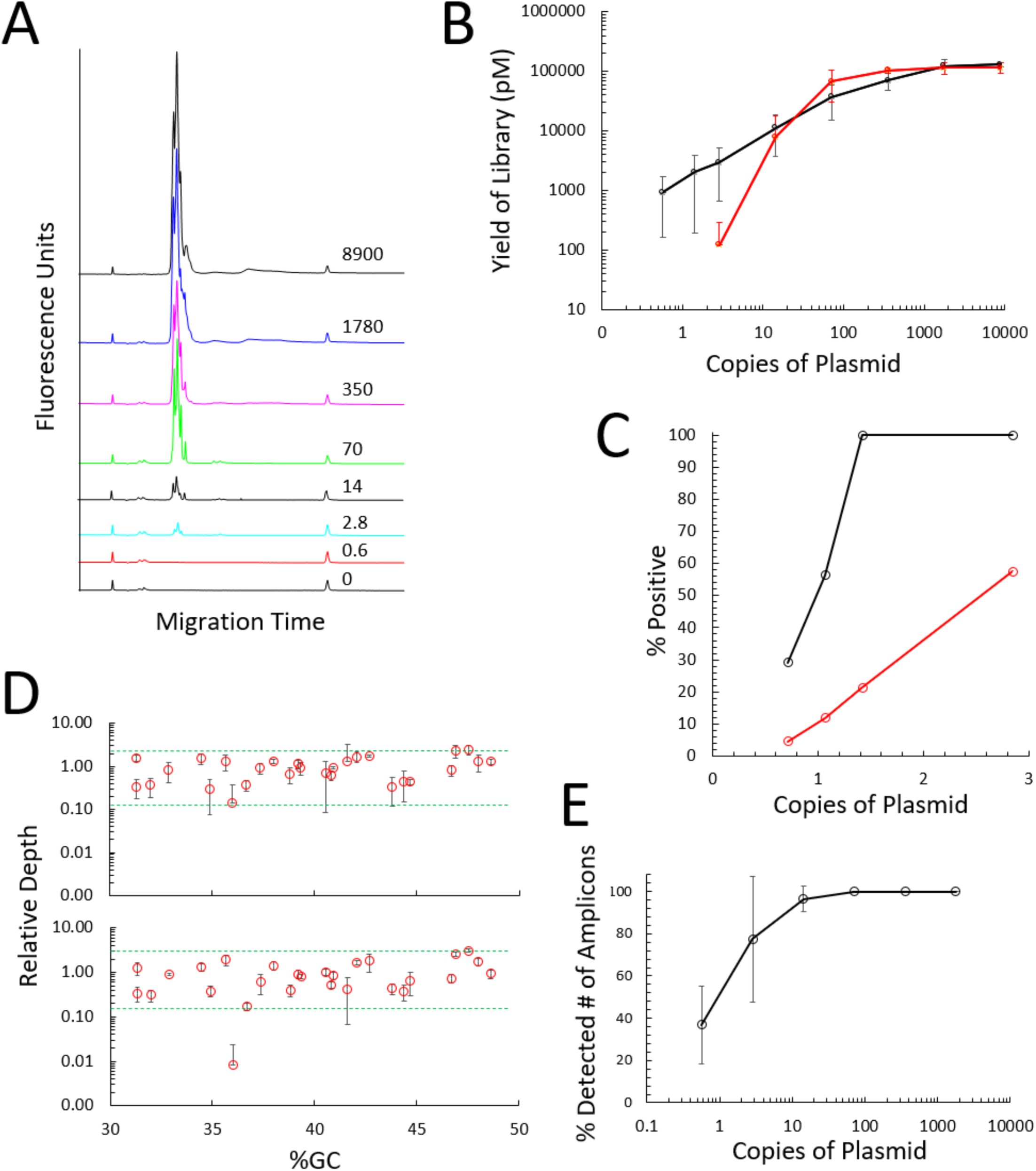
Detection of SARS-CoV-2 gene-containing plasmids by electrophoresis and sequencing. (**A**) Two plasmids, containing SARS-CoV-2 S and N genes, respectively, were diluted in human cDNA and amplified in multiplex PCR with pool 1 (172 pairs of primers). The number of plasmid copies per reaction, determined by ddPCR, were from 8,900 to 0.6. The resulting products obtained after multiplex PCR were resolved by electrophoresis. The specific amplification products (the library) can still be seen with 2.8 copies of plasmids. (**B**) The library yields can be detected down to 0.6 copies of plasmids (*n*=4) by multiplex PCR (black line), while only down to 2.8 copies by RT-PCR (> 4.5-fold difference) (red line). (**C**) Poisson process was used to estimate the chance of sampling around 1 copy of viral particles, and the mathematical model was used to estimate the chance of detecting them (red line). There is 12% of probability to detect 1.1 copies with a multiplex PCR efficiency of 6%. In reality, we observed a significantly higher 56% probability for 1.1 copies and 100% probability for 1.4 copies. (**D**) After sequencing 1.4 and 2.8 copies of plasmids, the reads of all 28 amplicons spanning both N and S genes were clustered within a 20-fold range of coverage (*n*=3). With 1.4 copies, only the reads of one amplicon were about 100-fold lower than the average (*n*=3). (**E**) About 96% of amplicons were recovered with 14 copies of plasmids, 77% with 2.8 copies, and 37% with 0.6 copies (*n*=3).

Estimated from the mathematical model described above, employing 28 amplicons provides a 16% chance to detect one single copy of the virus. We tested this predicted probability using one copy of plasmid in multiplex PCR reactions. The theoretical calculation gives a 66% probability to sample 1.1 copies, and a 12% chance to detect them based on a multiplex PCR efficiency of 6%. In practice, we experimentally observed a significantly higher 56% probability to detect 1.1 copies (Fig. 2C). These results suggest that the efficiency of multiplex PCR is actually higher than the previously estimated 6% when a single-stranded cDNA molecule is amplified.

When the amplified products were sequenced, we found that the recovered reads were within a range of about 20-fold relative depth with about 1.4 to 2.8 plasmids, and were uniformly distributed across the GC range (Fig. 2D). When detecting down to 1.4 copies of plasmids, only the reads from one amplicon were about 100-fold lower than the average. Approximately 96% of the amplicons were recovered with 14 copies of plasmids, 77% with 2.8 copies, and 37% with 0.6 copies (Fig. 2E).

### Detection of SARS-CoV-2 in COVID-19 specimens

The above two pools of primers were used to amplify the full SARS-CoV-2 genomes from a total of 13 nasopharyngeal swab specimens. These specimens were previously diagnosed to be SARS-CoV-2 positive by using RT-PCR. The viral load was found to be from 8 to 675,885 RNA copies/uL (Supplemental Table 2). These 13 viral genomes were successfully amplified and subsequently sequenced on the Illumina MiSeq. We found a correlation between genome coverage and viral copy number, as expected. While about 95% of the genome was covered at 100X for 8 copies of virus, 98-99% of the genomes were covered at 100X for 22 to 675,885 copies of virus at an average sequencing depth of 5,000 reads per amplicon (Fig. 3A). One genome, from the sample with 5,000 estimated viral copies, was covered at 96% for 100X. This coverage was lower than expected, and might have been caused by the poor sample quality resulting from processing or handling the viral RNA or library.

**Figure 3.**
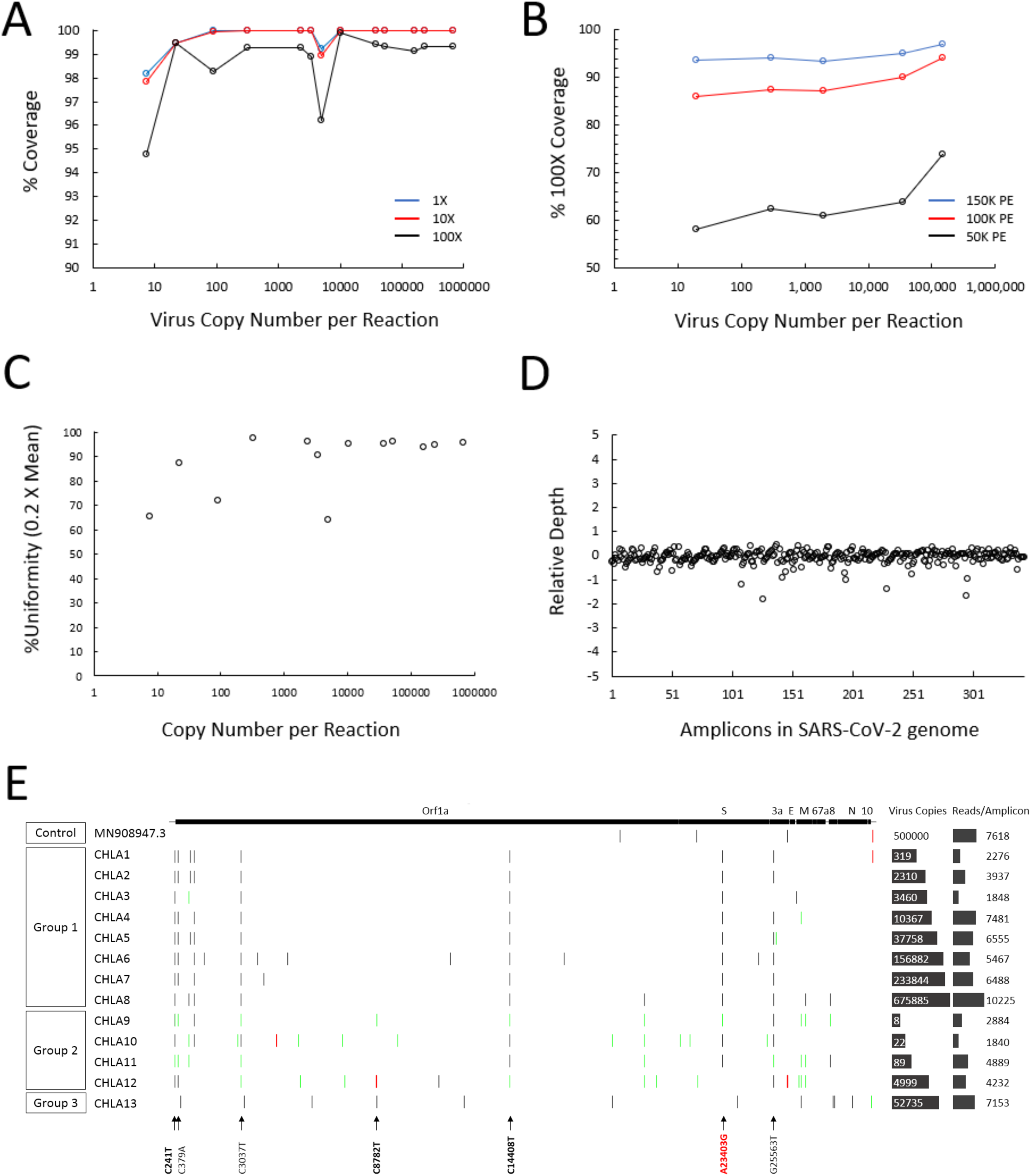
Sequencing the whole genome of SARS-CoV-2 in COVID-19 positive samples. (**A**). The entire genomes of SARS-CoV-2 were amplified and sequenced from a cohort of 13 COVID-19 positive patients. The copies of virus that were used in the multiplex PCR reaction range from 8 to 675,885. 98.3-99.9% of the genomes were covered at 100X with an average of 5,000 reads per amplicon from 22 to 675,885 copies. The coverage slightly decreased to 95% with 8 copies of virus. (**B**) Sub-sampling of 5 samples to 150,000 total reads slightly reduced the 100X coverage to 93-97%. (**C**) The amplification performance of the 343 amplicons for each sample was measured by the uniformity of 0.2X mean of the reads. Each circle represents one sample. (**D**) In one sample, the performance of each amplicon was evaluated by their log_10_ distance to the mean reads. Each circle represents one amplicon. (**E**). The mutations in each SARS-CoV-2 genome were detected and the SARS-CoV-2 samples were segregated into 3 groups based on similarities. The majority of the mutations were shared by groups 1 and 2. Group 2 contained low viral copy numbers in multiplex PCR, low reads per amplicon in sequencing and a considerable amount of apparent intra-host mutations. Mutations in group 3 were different from the other 2 groups. The mutations that associate with virulence^39^ are in bold. A23403G, which associates with high transmission^40^, is in red. The reference genome used was NC_045512.2. Black vertical lines represent point mutations, green vertical lines represent intra-host mutations, red vertical lines represent deletions.

CleanPlex libraries are usually sequenced at about 1,000 paired-end reads while still generating sufficient data for detecting mutations. To confirm SARS-CoV-2 libraries could be sequenced at about 1,000 reads per amplicon, we sub-sampled the data of 5 genomes to 150,000 and 100,000 total reads per library, which were equivalent to 437 and 291 reads per amplicon, respectively. At least 93% of the genome was covered at 100X with 150,000 total reads, and 86% of the genome was covered at 100X with 100,000 total reads. Even at 50,000 total reads per library, at least 58% of the genome was covered at 100X (Fig. 3B). The high coverage was also manifested by the superior uniformity of the number of amplicons amplified in the multiplex PCR reaction and recovered in the sequencing (Fig. 3C), and by the log_10_ distance of the number of each amplicon to the mean number of all amplicons in the library (Fig. 3D).

The mutations in the SARS-CoV-2 genome from the 13 specimens were detected by two independently developed algorithms. Only those mutations that were detected by both methods were reported. Assuming all viral particles from a single patient contained identical mutations, mutations with frequencies (%AF) > 60% were considered to be empirically true. The majority of the mutations identified in these 13 SARS-CoV-2 genomes clustered around 7 loci, probably reflecting the collection of these specimens in close communities or the transmission of the virus (Fig. 3E). According to the similarity of these mutations, samples were categorized into three groups. The majority of the mutations in the first group showed >98% mutation frequency. Groups 1 and 2 shared a considerable number of identical mutations, while Group 3 was significantly different, suggesting that the origin of this isolate might be traced back to a distinct lineage (Supplemental Table 3). Of note, all 13 strains contained at least one mutation that has been reported to be associated with SARS-CoV-2 virulence^39^. The D614G mutation (a G-to-A base change at position 23,403 in the reference strain NC_045512.2), which began spreading in Europe in early February, was then transmitted to new geographic regions and became a dominant form^40^, was found in 11 of these 13 specimens.

We considered mutations with %AF ≤ 60% as a likely reflection of intra-host heterogeneity^41^. To eliminate noise from PCR amplification and sequencing, only mutations with %AF ≥ 20% were considered. The 20% cutoff was selected based on the mutation profile from sequencing the synthetic SARS-CoV-2 RNA controls from Twist Bioscience. Some true intra-host mutations with %AF < 20% might be missed. Since no UMI was used in the multiplex PCR amplification, intra-host mutations might still be contaminated with noise originating from PCR amplification, even though a 20% cutoff was applied. The occurrence of such noise may be exacerbated by low viral copy inputs in the multiplex PCR, or low read depth per amplicon in sequencing. In groups 1 and 3, where both viral copy numbers and read depth per amplicon were high, only one intra-host mutation per genome was found in some of the specimens. In contrast, group 2 had low copy numbers, and low read depth per amplicon, and the numbers of apparent intra-host mutations were considerably higher (Fig. 3E and Supplemental Table 3). Some of these intra-host mutations occurred at the aforementioned 7 loci, or were found at other loci and were identical among the 4 specimens in group 2, while the remaining ones appeared random. Our findings suggested that these recurring mutations might not be true intra-host mutations. Indeed, it is possible that the low copy number inputs, as well as sequencing depth, caused a reduction in the %AF, and additionally introduced false intra-host mutations. Therefore, copy number and sequencing depth should be cautiously considered when a mutation is found to have low %AF.

### Metagenomic method design for novel pathogens

In order to characterize highly mutated viruses that would otherwise not be amplified by the pre-designed primer pairs, and to discover unknown pathogens, we subsequently developed a user-friendly multiplex-PCR-based metagenomic method. In this method, random hexamer-adapters were used to amplify DNA or cDNA targets in a multiplex PCR reaction. The large amounts of non-specific amplification products were removed by using Paragon Genomics’ background removal reagent, thus resolving a library suitable for sequencing. For RNA samples, Paragon Genomics’ reverse transcription reagents were used to convert RNA into cDNA, resulting in significantly reduced amount of human ribosomal RNA species.

We sequenced a library made with 4,500 copies of N and S gene-containing plasmids spiked into 10 ng of human gDNA, which roughly represents 3,300 haploid genomes. Even though the molar ratios of viral targets and human haploid genomes were comparable, the N and S genes, which encompass about 4kb of targets, were a negligible fraction of the 3 billion base pairs of a human genome. If every region of the human genome were amplified and sequenced at 0.6 million reads per sample, only one read of viral target would be recovered. In fact, our results showed that 16% of the recovered bases, or 13% of the recovered reads, were within the viral N and S genes (Fig. 4A and Supplemental Table 4). 80% of SARS-CoV-2 and 78% of mitochondrial targets were covered (Fig. 4B), and their base coverage was significantly higher than for human targets (Fig. 4C). In contrast, only 0.08% of human chromosomal regions were amplified. Furthermore, the human exonic regions were preferentially amplified (Fig. 4D). This suggested that the random hexamers deselected a large portion of the human genome, while favorably amplifying regions that were more “random” in base composition. Indeed, long gaps and lack of coverage in very large repetitive regions were observed in human chromosomes (Fig. 4E). On the contrary, the gaps in SARS-CoV-2 and mitochondrial regions were significantly shorter (Fig. 4F). We further optimized this method so that 96% of the SARS-CoV-2 genome was recovered (Fig. 4G). The depth of the recovered bases was within a 10-fold range on average. This 10-fold difference in coverage has been routinely observed with our multiplex PCR technology (Supplemental Fig. 4 and Supplemental Table 5). Therefore, increasing sequencing depth alone might not improve the coverage of the targeted regions further.

**Figure 4.**
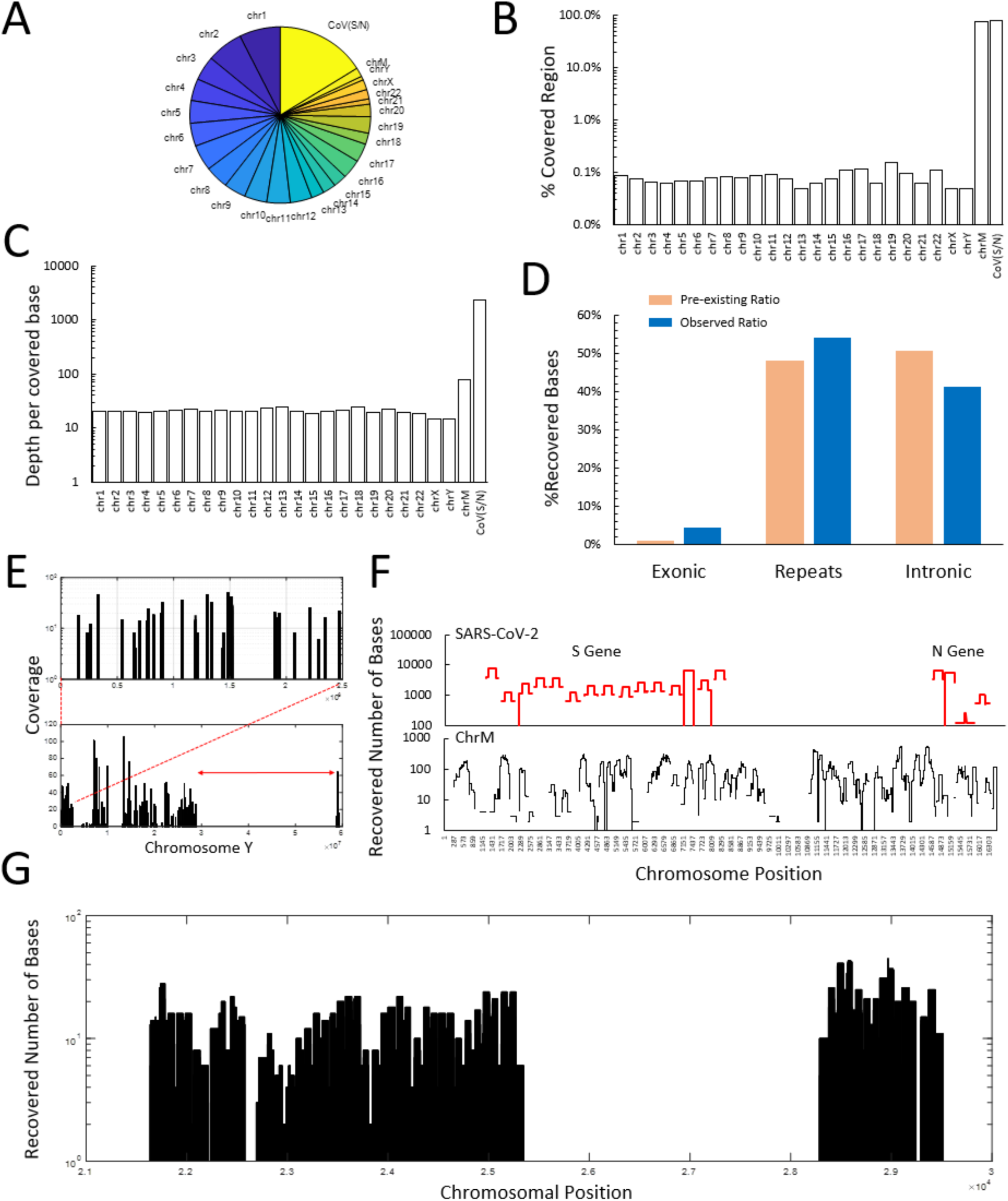
Multiplex PCR-based metagenomic method for the detection of SARS-CoV-2 genes. (**A**) Random hexamer-adapters were used in multiplex PCR to amplify 4,500 copies of plasmids in the background of 3,300 haploid human gDNA molecules. The resulting libraries were sequenced at an average of 0.6 million total reads. Of the total bases recovered, 16% were mapped to SARS-CoV-2 S and N genes. (**B**) 80% of S and N genes, and 78% of the human mitochondrial chromosome were amplified with >= 1X coverage, while only 0.08% of the human chromosomes were. (**C**) On average, S and N genes were covered at 2,346X, mitochondrial and human chromosomes at 77X and 20X, respectively. (**D**) Human exons were relatively over-amplified, at about 4-fold higher compared to their actual ratio within the genome. (**E**) Gaps and long regions of absence of amplification were observed for human chromosomes. An example shown here is chromosome Y. Small gaps were additionally found in the enlarged cluster of amplification. The long absent region (red double arrow) overlapped with the repetitive regions on Y chromosome. (**F**) Representation of the recovered regions in S and N genes and the human mitochondrial chromosome. The coverage was from 1,000- to 10,000-fold for S and N genes, and 30- to 500-fold for the mitochondrial chromosome. (**G**) This metagenomic method was subsequently improved, resulting in 95.7% of the SARS-CoV-2 S gene covered at 22.5X and 96.2% of the N gene covered at 12.8X, with a sequencing depth of 0.53M total reads.

## Discussion

This study provides a highly sensitive and robust multiplex PCR method for the detection of SARS-CoV-2. By amplifying hundreds of targets simultaneously, our multiplex PCR method is more sensitive than RT-PCR, and tolerates the presence of mutations in SARS-CoV-2. For the purpose of diagnosis, only one of two primer pools could be used, or the two pools could be alternatively used in adjacent samples to prevent cross contamination. While the amplification products from positive samples are mainly viral amplicons, low quantities of primer dimers are produced in the negative samples. Therefore, a simple measurement of the dsDNA concentration by fluorometry or spectrophotometry would not be sufficient. High-resolution electrophoresis is required to resolve the length of the amplification products in order to differentiate the target amplicons from the primer-dimers. Alternatively, a low-depth sequencing in the range of 50K reads per sample would provide definitive diagnostic results.

When both primer pools are used, the entire genome of SARS-CoV-2 can be enriched, sequenced and interrogated for the presence of any mutations. We demonstrated that mutations were detected from samples with viral loads ranging from 8 to half a million copies. For accurate sequencing and phylogenetic studies, a high-depth sequencing in the range of 300K reads per sample, along with an input of high viral load (>100 copies), are deemed necessary. We caution the interpretation of intra-host mutations obtained with a low input number of viral particles and in low sequencing depth data.

The current SARS-CoV-2 panel was demonstrated to specifically amplify the entire SARS-CoV-2 genome, and sequencing data obtained from the 13 COVID-19 RT-PCR positive samples clearly differentiate SARS-CoV-2 from other human coronaviruses, such as MERS-CoV, CoV 229E, CoV OC43, CoV NL63, CoV HKU1. 0% of the obtained sequencing reads from each of the 13 samples were aligned to the genome sequence of any of the above viral species. This is in contrast to what was reported for a similar SARS-CoV-2 panel^42^. Such high specificity argues strongly that this panel could be further expanded to include simultaneous detection of other respiratory viruses including influenza virus.

Metagenomic method is a powerful technology that can theoretically detect any sequences in the specimens. However, metagenomic methods usually require very high sequencing depth in order to find the target sequences, and hence are economically prohibitive as a diagnostic assay. To overcome this constraint, we developed a multiplex-PCR-based metagenomic method that achieved >96% coverage of the S and N genes of SARS-CoV-2 in the contest of human gDNA, while only required ∼0.6M of total reads per library. This coverage was superior given the recommended 50% threshold of coverage for drafting a genome^43^. The results were obtained with no additional means of host depletion to remove human gDNA and rRNA. The viral bases were 16% of the total recovered bases in the sequencing. Yet it still necessary to verify and validate the detection of SARS-CoV-2 and the other coronaviruses and respiratory viruses by this metagenomic method.

## Materials and Methods

### Ethics statement

Clinical SARS-CoV-2 samples were collected at Children’s Hospital of Los Angeles. Samples and ancillary clinical and epidemiological data were de-identified before analysis, and are thus considered exempt from human subject regulations, with a waiver of informed consent according to 45 CFR 46.101(b) of the US Department of Health and Human Services. Analysis of the nasopharyngeal swab samples from patients with COVID-19 disease was approved by the Ministry of Health in the US. Patients in the 2020 COVID-19 outbreak from 1 January 2020 to 30 August 2020 provided oral consent for study enrolment and the collection and analysis of their nasopharyngeal swab. Consent was obtained at the homes of patients or in hospital isolation wards by a team that included staff members of Children’s Hospital of Los Angeles. SARS-CoV-2 viruses were purified from the clinical samples by using QIAamp Viral RNA Mini Kit (Qiagen, Cat. No. 52906). The preparations were analyzed by real-time RT–PCR testing for the determination of viral titers of SARS-CoV-2 by standard curve analysis. The full genomes of SARS-CoV-2 viruses were amplified by using the CleanPlex SARS-CoV-2 panel (Paragon Genomics, SKU 918011) and sequenced on an Illumina MiSeq at Children’s Hospital of Los Angeles.

### Materials

The Universal Human Reference RNA was from Agilent Technologies, Inc. (Cat#74000). The plasmids containing either S or N gene of SARS-CoV-2 (pUC-S and pUC-N, respectively) were purchased from Sangon Biotech, Shanghai, China. The PCR primers used in ddPCR and RT-PCR reactions for S gene are 5’-TGTACTTGGACAATCAAAAAGAGTTGAT and 5’-AGGAGCAGTTGTGAAGTTCTTTTC; for N gene are 5’-GGGGAACTTCTCCTGCTAGAAT and 5’-CAGACATTTTGCTCTCAAGCTG, respectively.

### Multiplex PCR panel design

Panel design is based on the SARS-CoV-2 sequence NC_045512.2 (https://www.ncbi.nlm.nih.gov/nuccore/NC_045512.2/). In total, 343 primer pairs, distributed into two separate pools, were selected by a proprietary panel design pipeline to cover the whole viral genome except for 92 bases at its ends. Primers were optimized to preferentially amplify the SARS-CoV-2 cDNA versus background human cDNA or genomic DNA. They were also optimized to amplify the covered genome uniformly.

### Reverse transcription

50ng of Universal Human Reference RNA was converted into cDNA using random primers and SuperScript IV Reverse Transcriptase, following the supplier recommended method (Thermo Fisher Scientific, Cat# 18090050). After reverse transcription, cDNA was purified with 2.4X volume of magnetic beads, and washed twice with 70% ethanol. Finally, the purified cDNA was dissolved in 1X TE buffer and used per multiplex PCR reaction.

### RT-PCR

Plasmids pUC-S and pUC-N, in combination with human cDNA, were used in each reaction. Paragon Genomics’ CleanPlex secondary PCR mix was used with 100nM of each PCR primers in 10ul reactions. The PCR thermal cycling protocol used was 95°C for 10min, then 98°C for 15sec, 60°C for 30sec for 45 cycles.

### ddPCR

ddPCR was performed on QX200 from Bio-Rad. Plasmids pUC-S and pUC-N at the estimated copy numbers 1 (6 repeats), 2 (3 repeats), and 100 (3 repeats) were tested. In each reaction, the ddPCR thermal cycling protocol used was 95°C for 5min, then 95°C for 30sec, 60°C for 1min with 60 cycles, 4°C for 5min and 90°C for 5min, 4°C hold. The resulting data were analyzed by following the supplier recommended method.

### Multiplex PCR

Paragon Genomics’ CleanPlex multiplex PCR reagents and protocol were used. Briefly, a 10µl multiplex PCR reaction was made by combining 5X mPCR mix, 10X Pool 1 of the panel, water and viral template cDNA. The reaction was run in a thermal cycler (95°C for 10min, then 98°C for 15sec, 60°C for 5min for 10 cycles), then terminated by the addition of 2µl of stop buffer. The reaction was then purified by 29µl of magnetic beads, followed by a secondary PCR with a pair of primers for 25 cycles. The secondary PCR added sample indexes and sequencing adapters, allowing for sequencing of the resulting products by high throughput sequencing. A final bead purification was performed after the secondary PCR, followed by library interrogation using a Bioanalyzer 2100 instrument with Agilent High Sensitivity DNA Kit (Agilent Technologies, Inc. Part# 5067-4626).

### Mathematical Modeling

A cumulative Poisson probability was used to build the mathematical model. In Microsoft Excel, the following function was used:

*P* = 1 - POISSON.DIST(1, ***λ***, TRUE)

For multiplex PCR, ***λ*** = *f* × 80% × *n* × *m*, where *f* = frequency of target(s) per genome. *f* = *l*/*L*, where *l* = cumulative length of amplicon, *L* = length of genome. For a panel of 172 amplicons, *l* = 172 × average length of amplicon. *n* = number of virus genomes in the sample (*n* = 1,2,3…); *m* = amplification efficiency. 80% of targets were assumed to be successfully amplified in multiplex PCR. For RT-PCR, ***λ*** = *f* × *n* × *m. q* = number of detected virus genomes. *q* is used to plot the graph reported in the paper, e.g., the copies of virus against the matching probabilities. For multiplex PCR, *q* = *n*/*a, a* = the number of amplicons used in the multiplex PCR. For RT-PCR, *q* = *n* × *f*.

### Multiplex-PCR-based metagenomic method

Paragon Genomics’ CleanPlex metagenomic reagents and protocol were used. Briefly, a 10µl multiplex PCR reaction was made by combining 5X mPCR mix, 10X random hexamer-adapters, water and the viral template cDNA. The PCR thermal cycling protocol used was 95°C for 10min, then 98°C for 15sec, 25°C for 2min, 60°C for 5min for 10 cycles. The reaction was then terminated by the addition of 2µl of stop buffer, and purified by 29µl of magnetic beads. The resulting solution was treated with 2µl of CleanPlex reagent at 37°C for 10min to remove non-specific amplification products. After a magnetic bead purification, the product was further amplified in a secondary PCR with a pair of primers for 25 cycles to produce the metagenomic library. This metagenomic library was further purified by magnetic beads before sequencing.

### High throughput NGS sequencing and data analysis

High throughput NGS sequencing was performed using Illumina iSeq 100, MiSeq and MGI sequencers (DNBSEQ-G400 and its research-grade CoolMPS sequencing kits). Detailed information for the samples sequenced and used in this manuscript can be found in Supplemental Table 4. Raw sequencing data were trimmed for adaptors using cutadapt version 1.14. The sequences obtained were mapped to the SARS-CoV-2 genome (NC_045512.2) with bwa-mem using Sentieon version 201808.01. Duplicate read marking was skipped. Base-quality recalibration, re-alignment of indels and quality metrics was accomplished with Sentieon. The resulting BAM files were then used to calculate depth and coverage metrics using Samtools version 1.3.1. Algorithms developed independently at Children’s Hospital of Los Angeles and paragon Genomics were sued to detect the mutations in the genome of SARS-CoV-2.

## Data availability

All sequencing data used in this publication are available for downloading at NCBI’s Sequence Read Archive (https://www.ncbi.nlm.nih.gov/Traces/study/?acc=PRJNA614546).

## Acknowledgements

The authors would like to thank Jian Xu of MGI, BGI-Shenzhen for technical support in MGI sequencing, Dr. Jin Billy Li of Stanford University for the coordination of academic support, Dr. Alexander E. Urban of Stanford University for suggestions in preparing the manuscript, and Dr. Zihuai He of Stanford University for discussions on a potential mathematical model of the metagenomic method.

## Author contributions

C.L., D.N.D., Z.L. conceived the study and drafted the manuscript. C.L., D.N.D., J.S., V.K., L.Y.L., L.L., R.F., G.B., J.L., A.O., G.L., Z.L. performed experiments, analysis and revised the manuscript. U. P. performed the initial diagnosis and quantification of clinical COVID-19 samples. D. O and D. R. performed CleanPlex library preparation and NGS sequencing of the positive COVID-19 clinical samples with MiSeq. M. B., D. M., A. R., and L. S. analyzed the data from the clinical samples. X. G. and J. D. B. supervised the study of clinical samples and revised the manuscript. B.Z., A.E.U. performed ddPCR experiments and analysis. A.B., H.T. performed MGI sequencing and analysis.

## Conflict of Interest

The authors declare no competing interests.

## Supplemental Information

**Supplemental Fig 1.**
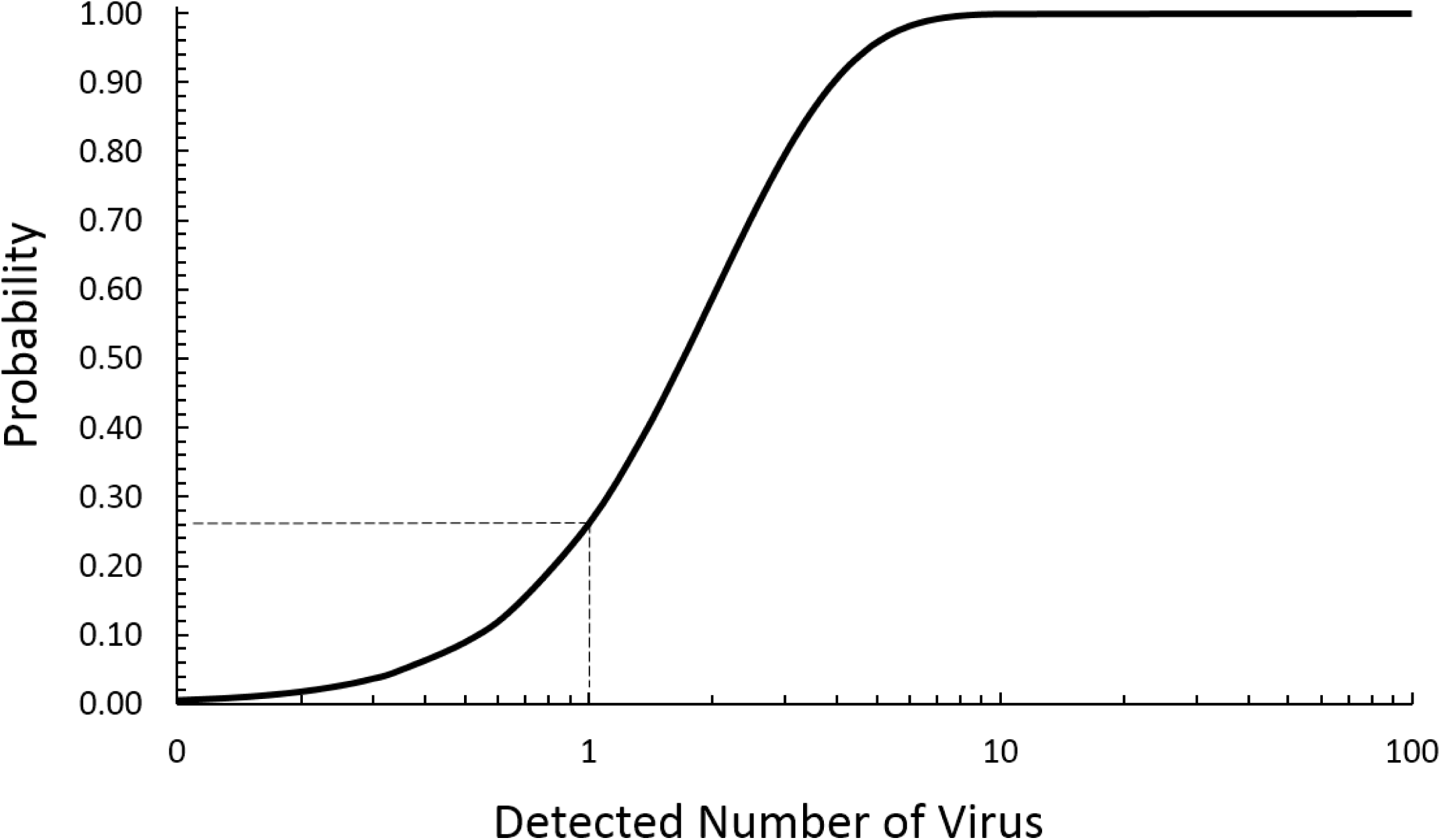
A mathematical model of RT-PCR. The same model was used to estimate the LOD of both RT-PCR and multiplex PCR, through changing the amplicon length and number, the virus genome size, as well as the intended detected copies and PCR efficiency. We found that the probability of detecting 1 copy of SARS-CoV-2 is 26% by using RT-PCR, and the LOD is independent of the length of virus genome.

**Supplemental Fig 2.**
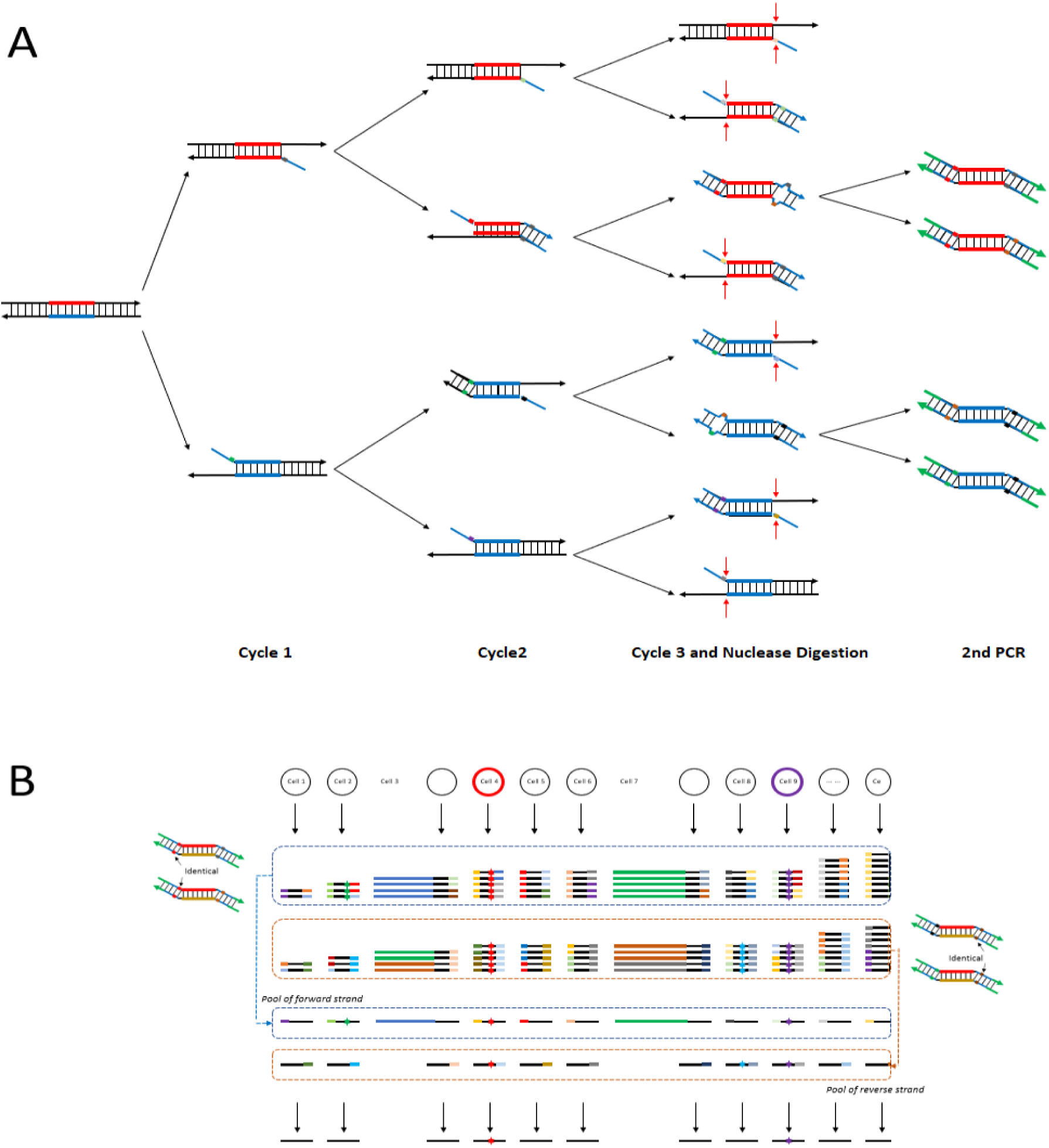
Multiplex PCR efficiency as determined by using CleanPlex^®^ UMI technology by Paragon Genomics. **A.** The underlying mechanism of CleanPlex^®^ UMI technology by Paragon Genomics, which uses three cycles of multiplex PCR to label targets with UMI. The redundant UMIs generated in the third cycle of PCR are removed through nuclease digestion of the single-stranded regions. The resulting products are further amplified by using a pair of universal primers, while sample indexes and sequencing adapters are introduced. **B.** The UMIs are initially sorted based on the UMI itself, and further on the occurrence of identical UMIs on either the 5′ or 3′ end of amplicons after sequencing, thus allowing the identification of the original template. The position of identical UMIs on either the 5′ or 3′ end of amplicons can further indicate whether the final amplification products are from the pool of the sense or antisense strand of the original templates.

**Supplemental Fig 3.**
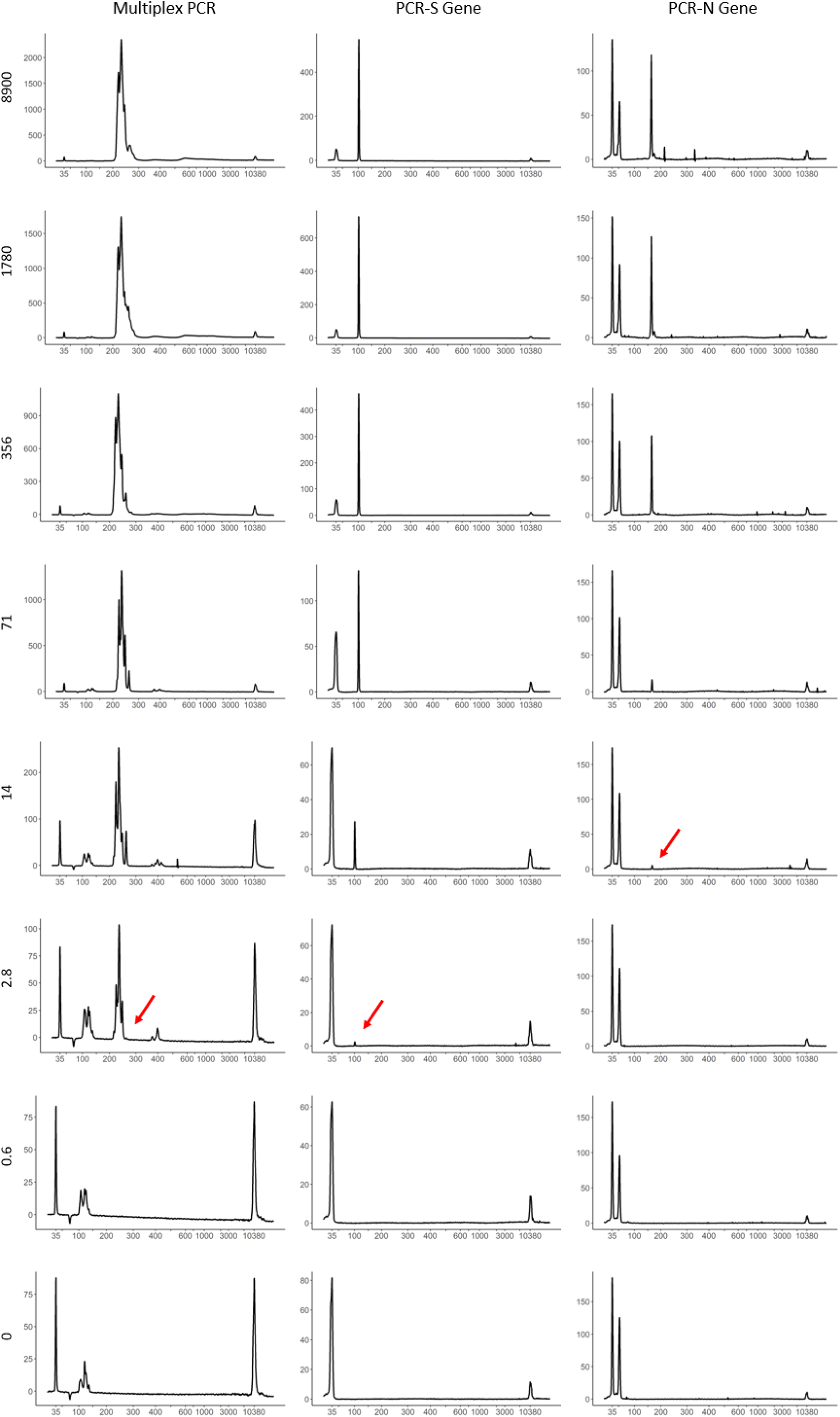
Comparison of LOD between multiplex PCR and regular PCR. A total of 35 cycles was used in multiplex PCR, while 45 cycles were used for regular PCR. The resulting amplification products from multiplex PCR were processed as described in the Materials and Methods. The PCR products were directly resolved using high sensitivity DNA chips on a Bioanalyzer 2100 instrument. X-axis indicates fragment size (bp) and y-axis indicates fluorescence units. The arrows point to the expected specific amplification products. The number of plasmid copies is indicated on the left.

**Supplemental Fig 4.**
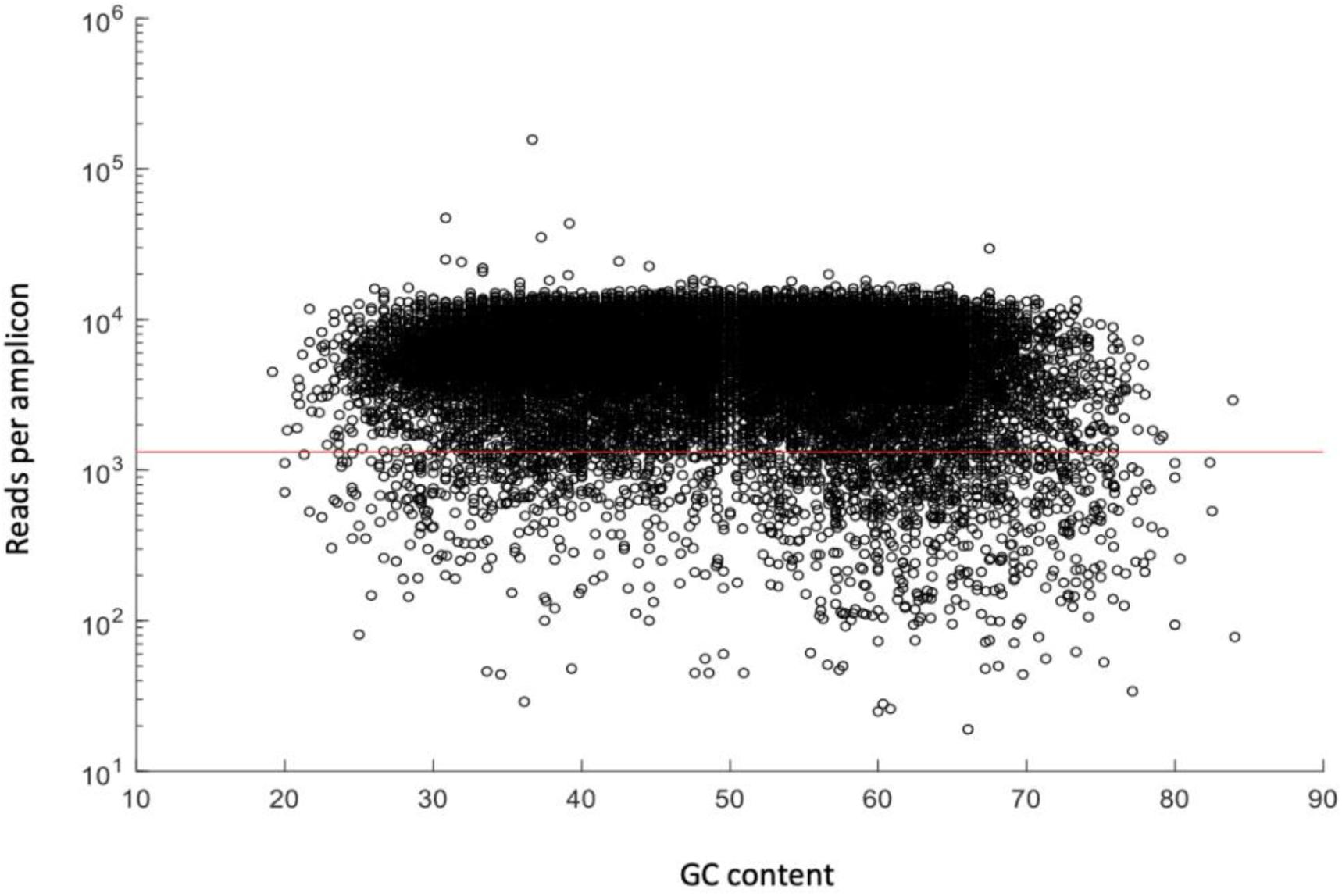
Performance statistics of the amplicons retrieved from multiplex PCR method highlighting a 10-fold range read depth. The number of sequencing reads for a majority of the recovered amplicons (Supplemental Table 5) were within a 10-fold range, representing a uniformity of 92.62 ± 1.96% at 0.2X mean (red line).

**Supplemental Table 1.**
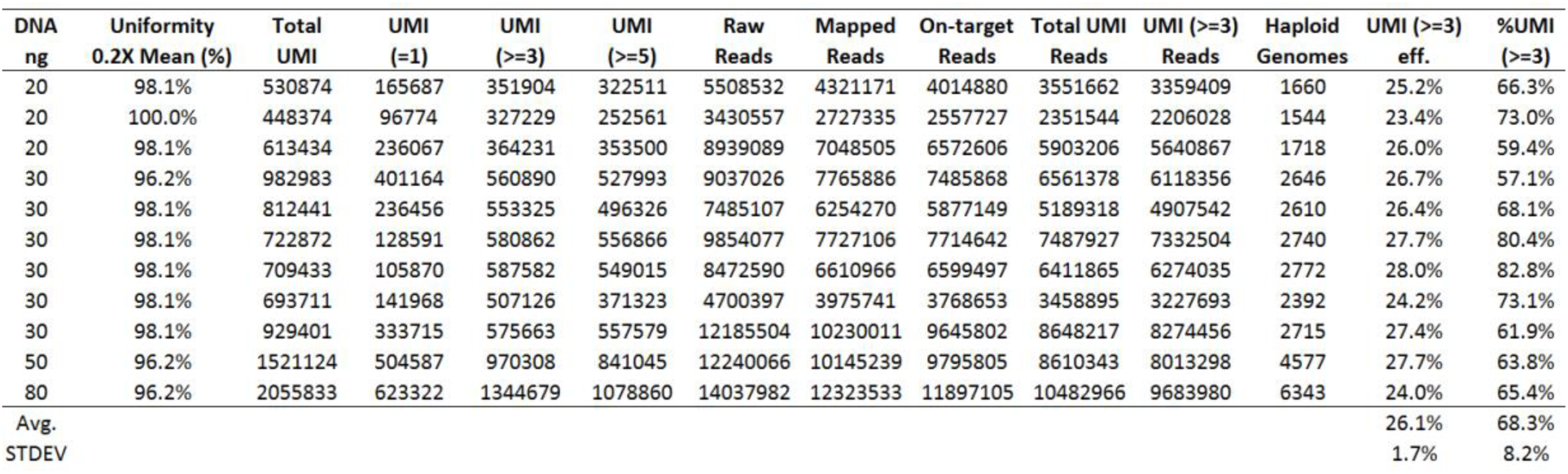
Multiplex PCR efficiency as determined by using CleanPlex^®^ UMI technology by Paragon Genomics. A 53-amplicon panel was amplified with 20 to 80 ng of gDNA (NA12878) by using CleanPlex^®^ UMI technology by Paragon Genomics, the mechanism of which is depicted in Supplemental Fig 2. The efficiency of multiplex PCR was found to be 26%, calculated from the recovered numbers of UMI clusters that contained >= 3 members (UMI (>=3)).

**Supplemental Table 2.** RT-PCR and sequencing results of SARS-CoV-2 genomes amplified from a cohort of 13 COVID-19 positive patients.

| ID | Copy Number | Uniformity<br>0.2X Mean (%) | Mapping<br>Rate (%) | On-Target<br>Rate (%) | Total<br>Reads | Mapped<br>Reads | On-Target<br>Reads | Primer<br>Dimer Reads | Primer Dimer<br>Rate (%) | Average On-Target Reads<br>per Amplicon |
| --- | --- | --- | --- | --- | --- | --- | --- | --- | --- | --- |
| CHLA9 | 8 | 65.31 | 99.81 | 89.26 | 1110606 | 1108547 | 989515 | 493 | 0.05 | 2884 |
| CHLA10 | 22 | 87.46 | 84.51 | 93.14 | 845394 | 714470 | 665443 | 20129 | 3.02 | 1940 |
| CHLA11 | 89 | 71.72 | 99.90 | 96.47 | 1740180 | 1738420 | 1677034 | 869 | 0.05 | 4889 |
| CHLA1 | 319 | 97.38 | 98.03 | 99.21 | 802986 | 787187 | 780950 | 1894 | 0.24 | 2276 |
| CHLA2 | 2310 | 96.21 | 95.92 | 99.13 | 1420132 | 1362254 | 1350437 | 7013 | 0.52 | 3937 |
| CHLA3 | 3460 | 90.67 | 96.8 | 98.42 | 665536 | 644217 | 634048 | 3132 | 0.49 | 1848 |
| CHLA12 | 4999 | 63.85 | 99.73 | 93.83 | 1551346 | 1547177 | 1451691 | 8812 | 0.61 | 4232 |
| CHLA4 | 10367 | 95.04 | 99.83 | 98.08 | 2620898 | 2616444 | 2566262 | 1585 | 0.06 | 7481 |
| CHLA5 | 37758 | 95.04 | 99.32 | 99.20 | 2282126 | 2266649 | 2248408 | 1235 | 0.05 | 6555 |
| CHLA13 | 52735 | 95.92 | 99.94 | 98.96 | 2480938 | 2479370 | 2453510 | 186 | 0.01 | 7153 |
| CHLA6 | 156882 | 93.59 | 99.77 | 99.43 | 1890384 | 1885967 | 1875238 | 462 | 0.02 | 5467 |
| CHLA7 | 233844 | 94.75 | 99.79 | 99.00 | 2252828 | 2248089 | 2225503 | 308 | 0.01 | 6488 |
| CHLA8 | 675855 | 95.63 | 99.96 | 97.63 | 3594226 | 3592758 | 3507470 | 49 | 0.00 | 10225 |
| MN908947.3 | 500000 | 90.96 | 99.71 | 98.64 | 2656772 | 2649047 | 2613012 | 271 | 0.01 | 7618 |

**Supplemental Table 3.**
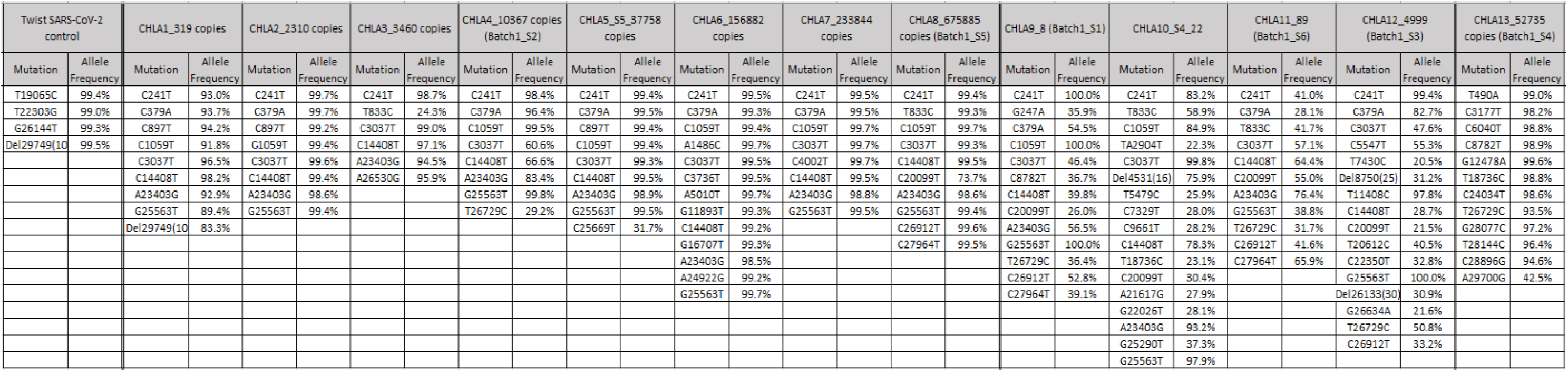
Detected SARS-CoV-2 mutations in a cohort of 13 COVID-19 positive patients.

**Supplemental Table 4.**
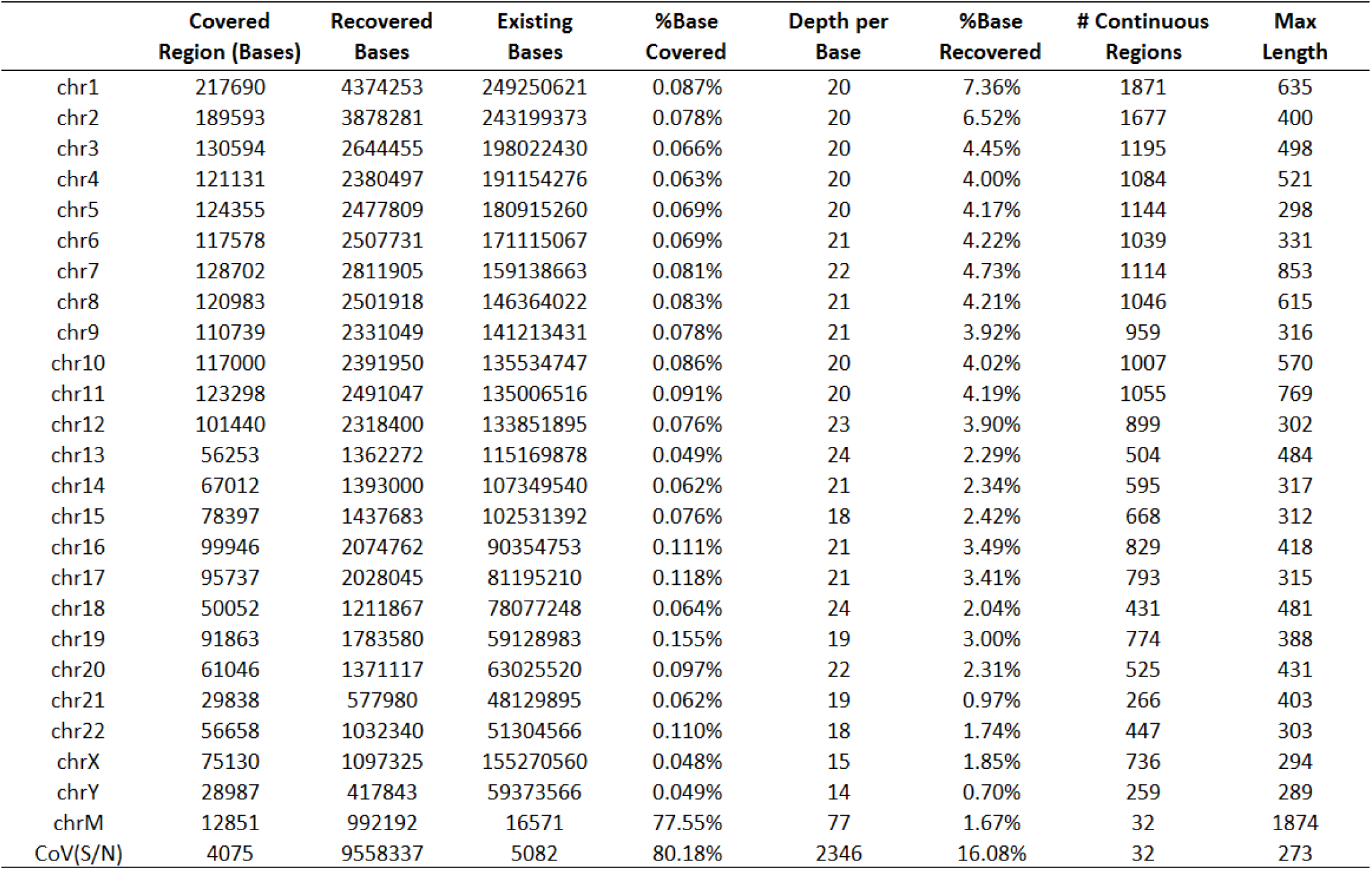
Sequencing results of the multiplex PCR-based metagenomic method using 4,500 copies of plasmids containing SARS-CoV-2 S and N genes spiked in 10ng of human gDNA.

|  | Covered<br>Region (Bases) | Recovered<br>Bases | Existing<br>Bases | %Base<br>Covered | Depth per<br>Base | %Base<br>Recovered | # Continuous<br>Regions | Max<br>Length |
| --- | --- | --- | --- | --- | --- | --- | --- | --- |
| chr1 | 217690 | 4374253 | 249250621 | 0.087% | 20 | 7.36% | 1871 | 635 |
| chr2 | 189593 | 3878281 | 243199373 | 0.078% | 20 | 6.52% | 1677 | 400 |
| chr3 | 130594 | 2644455 | 198022430 | 0.066% | 20 | 4.45% | 1195 | 498 |
| chr4 | 121131 | 2380497 | 191154276 | 0.063% | 20 | 4.00% | 1084 | 521 |
| chr5 | 124355 | 2477809 | 180915260 | 0.069% | 20 | 4.17% | 1144 | 298 |
| chr6 | 117578 | 2507731 | 171115067 | 0.069% | 21 | 4.22% | 1039 | 331 |
| chr7 | 128702 | 2811905 | 159138663 | 0.081% | 22 | 4.73% | 1114 | 853 |
| chr8 | 120983 | 2501918 | 146364022 | 0.083% | 21 | 4.21% | 1046 | 615 |
| chr9 | 110739 | 2331049 | 141213431 | 0.078% | 21 | 3.92% | 959 | 316 |
| chr10 | 117000 | 2391950 | 135534747 | 0.086% | 20 | 4.02% | 1007 | 570 |
| chr11 | 123298 | 2491047 | 135006516 | 0.091% | 20 | 4.19% | 1055 | 769 |
| chr12 | 101440 | 2318400 | 133851895 | 0.076% | 23 | 3.90% | 899 | 302 |
| chr13 | 56253 | 1362272 | 115169878 | 0.049% | 24 | 2.29% | 504 | 484 |
| chr14 | 67012 | 1393000 | 107349540 | 0.062% | 21 | 2.34% | 595 | 317 |
| chr15 | 78397 | 1437683 | 102531392 | 0.076% | 18 | 2.42% | 668 | 312 |
| chr16 | 99946 | 2074762 | 90354753 | 0.111% | 21 | 3.49% | 829 | 418 |
| chr17 | 95737 | 2028045 | 81195210 | 0.118% | 21 | 3.41% | 793 | 315 |
| chr18 | 50052 | 1211867 | 78077248 | 0.064% | 24 | 2.04% | 431 | 481 |
| chr19 | 91863 | 1783580 | 59128983 | 0.155% | 19 | 3.00% | 774 | 388 |
| chr20 | 61046 | 1371117 | 63025520 | 0.097% | 22 | 2.31% | 525 | 431 |
| chr21 | 29838 | 577980 | 48129895 | 0.062% | 19 | 0.97% | 266 | 403 |
| chr22 | 56658 | 1032340 | 51304566 | 0.110% | 18 | 1.74% | 447 | 303 |
| chrX | 75130 | 1097325 | 155270560 | 0.048% | 15 | 1.85% | 736 | 294 |
| chrY | 28987 | 417843 | 59373566 | 0.049% | 14 | 0.70% | 259 | 289 |
| chrM | 12851 | 992192 | 16571 | 77.55% | 77 | 1.67% | 32 | 1874 |
| CoV(S/N) | 4075 | 9558337 | 5082 | 80.18% | 2346 | 16.08% | 32 | 273 |

**Supplemental Table 5.** Performance statistics of the amplicons retrieved from our multiplex PCR method highlighting a 10-fold range of read depth. To simulate multiplex PCR with random hexamers as primers, we used a panel of 27,296 pairs of primers to perform multiplex PCR. These primers were divided into 2 overlapping primer pools, and amplification was initially performed in two separate reactions. The number of sequencing reads for a majority of the recovered amplicons were within a 10-fold range, representing a uniformity of 92.62 ± 1.96% at 0.2X mean.

| Sample Number | Uniformity 0.2X Mean (%) | Mapping Rate (%) | On-Target Rate (%) | Total Reads | Mapped Reads | On-Target Reads | Primer Dimer Reads | Primer Dimer Rate (%) | Average On-Target Reads per Amplicon |
| --- | --- | --- | --- | --- | --- | --- | --- | --- | --- |
| 1 | 93.6% | 94.0% | 96.2% | 269435076 | 253341610 | 243602321 | 3326193 | 1.31% | 8924 |
| 2 | 92.7% | 97.3% | 95.9% | 272733262 | 265270282 | 254306010 | 1920151 | 0.72% | 9316 |
| 3 | 92.8% | 96.6% | 95.8% | 188776124 | 182357219 | 174778001 | 1492049 | 0.82% | 6403 |
| 4 | 93.1% | 97.2% | 95.7% | 265645114 | 258182429 | 247093224 | 1879080 | 0.73% | 9052 |
| 5 | 92.6% | 97.1% | 95.5% | 232875020 | 226046731 | 215875589 | 1535441 | 0.68% | 7908 |
| 6 | 93.5% | 97.1% | 96.2% | 192403072 | 186821637 | 179683779 | 1266338 | 0.68% | 6582 |
| 7 | 91.3% | 96.5% | 96.5% | 181465458 | 175141658 | 168975648 | 1423714 | 0.81% | 6190 |
| 8 | 88.1% | 96.7% | 96.6% | 191995346 | 185607975 | 179350573 | 1390209 | 0.75% | 6570 |
| 9 | 95.0% | 97.2% | 95.3% | 197356014 | 191797447 | 182683756 | 1056748 | 0.55% | 6692 |
| 10 | 95.0% | 96.5% | 95.7% | 201474266 | 194391114 | 185997107 | 1317629 | 0.68% | 6814 |
| 11 | 95.0% | 96.7% | 96.5% | 193125370 | 186751406 | 180154727 | 1036439 | 0.55% | 6600 |
| 12 | 88.2% | 95.5% | 97.0% | 199616350 | 190581763 | 184843808 | 1908098 | 1.00% | 6771 |
| 13 | 92.0% | 93.3% | 96.4% | 272873692 | 254679030 | 245480306 | 3824568 | 1.50% | 8993 |
| 14 | 94.0% | 93.6% | 95.9% | 297541418 | 278622642 | 267164257 | 4259698 | 1.53% | 9787 |
| 15 | 91.2% | 91.7% | 96.3% | 323595774 | 296801010 | 285698174 | 5184019 | 1.75% | 10466 |
| 16 | 91.4% | 93.9% | 96.3% | 248888684 | 233638153 | 224968924 | 5337352 | 2.28% | 8241 |
| 17 | 91.5% | 97.3% | 96.8% | 206702726 | 201183261 | 194776826 | 2063586 | 1.03% | 7135 |
| 18 | 91.5% | 98.2% | 96.9% | 178948198 | 175675252 | 170297061 | 1083447 | 0.62% | 6238 |
| 19 | 90.2% | 99.0% | 97.1% | 206684484 | 204582941 | 198563161 | 708813 | 0.35% | 7274 |
| 20 | 91.1% | 87.3% | 95.0% | 397262166 | 346621107 | 329334881 | 17048174 | 4.92% | 12065 |
| 21 | 94.5% | 95.5% | 96.1% | 215357334 | 205717503 | 197695646 | 2174832 | 1.06% | 7242 |
| 22 | 94.1% | 99.1% | 95.9% | 166942258 | 165384936 | 158665846 | 425846 | 0.26% | 5812 |
| 23 | 94.2% | 98.7% | 96.0% | 198067370 | 195451983 | 187701462 | 631609 | 0.32% | 6876 |
| 24 | 94.2% | 98.6% | 96.1% | 169536088 | 167192888 | 160657429 | 514940 | 0.31% | 5885 |
| 25 | 94.8% | 91.4% | 94.9% | 356335780 | 325681114 | 308973305 | 8034601 | 2.47% | 11319 |
| Avg. | 92.6% | 95.8% | 96.1% |  |  |  |  | 1.11% | 7806 |
| STDEV | 1.96% | 2.77% | 0.58% |  |  |  |  | 0.98% | 1739 |
| CV | 2.12% | 2.89% | 0.60% |  |  |  |  | 88.6% | 22.3% |

**Supplemental Table 6.** Detailed information for the 45 sequenced samples used in this manuscript. NCBI’s Sequence Read Archive (SRA) Sample ID and accession numbers for all 45 sequenced samples, along with sequencing details, sample description and the figure in which each sample’s data has been used is listed in this table. These fastq files are available for downloading at https://www.ncbi.nlm.nih.gov/Traces/study/?acc=PRJNA614546

| Sample | File Name | Figure | Content | Accession | Isolation Source |
| --- | --- | --- | --- | --- | --- |
| 10 | ILLUMINA (Illumina iSeq 100) run: 193,681 spots, 58.5M bases, 15.6Mb downloads | Figure 2D | 2.8 copies_1 | SRX7976138 | Plasmid |
| 9 | ILLUMINA (Illumina iSeq 100) run: 222,876 spots, 67.3M bases, 17.9Mb downloads | Figure 2D | 2.8 copies_2 | SRX7976139 | Plasmid |
| 8 | ILLUMINA (Illumina iSeq 100) run: 18,392 spots, 5.6M bases, 1.7Mb downloads | Figure 2D | 2.8 copies_3 | SRX7976140 | Plasmid |
| 7 | ILLUMINA (Illumina iSeq 100) run: 900,360 spots, 271.9M bases, 72.1Mb downloads | Figure 2D | 1.4 copies_1 | SRX7976141 | Plasmid |
| 6 | ILLUMINA (Illumina iSeq 100) run: 144,240 spots, 43.6M bases, 11.8Mb downloads | Figure 2D | 1.4 copies_2 | SRX7976142 | Plasmid |
| 5 | ILLUMINA (Illumina iSeq 100) run: 977,584 spots, 295.2M bases, 83.5Mb downloads | Figure 2D | 1.4 copies_3 | SRX7976143 | Plasmid |
| 4 | ILLUMINA (Illumina iSeq 100) run: 457,181 spots, 138.1M bases, 46.9Mb downloads | Figure 3 | Metagenomics | SRX7976144 | Plasmid |
| 3 | ILLUMINA (Illumina iSeq 100) run: 517,641 spots, 156.3M bases, 45Mb downloads | Figure 4 | Meta_20 dilution | SRX7976145 | Plasmid |
| 2 | ILLUMINA (Illumina iSeq 100) run: 613,188 spots, 185.2M bases, 53.5Mb downloads | Figure 4 | Meta_4 dilution | SRX7976146 | Plasmid |
| 1 | ILLUMINA (Illumina iSeq 100) run: 565,052 spots, 170.6M bases, 49.2Mb downloads | Figure 4 | Meta_0.8 dilution | SRX7976147 | Plasmid |
| 31 | BGISEQ (BGISEQ-500) run: 21M spots, 2.1G bases, 1Gb downloads | Figure 2E | 500 dilution_1 | SRX7990909 | Plasmid |
| 30 | BGISEQ (BGISEQ-500) run: 21.5M spots, 2.1G bases, 1Gb downloads | Figure 2E | 100 dilution_1 | SRX7990910 | Plasmid |
| 17 | BGISEQ (BGISEQ-500) run: 16.5M spots, 1.7G bases, 817.7Mb downloads | Figure 2E | 20 dilution_1 | SRX7990923 | Plasmid |
| 16 | BGISEQ (BGISEQ-500) run: 18.4M spots, 1.8G bases, 917.7Mb downloads | Figure 2E | 4 dilution_1 | SRX7990924 | Plasmid |
| 15 | BGISEQ (BGISEQ-500) run: 24.7M spots, 2.5G bases, 1.2Gb downloads | Figure 2E | 0.8 dilution_1 | SRX7990925 | Plasmid |
| 14 | BGISEQ (BGISEQ-500) run: 9.2M spots, 924.4M bases, 420.4Mb downloads | Figure 2E | 0.16 dilution_1 | SRX7990926 | Plasmid |
| 13 | BGISEQ (BGISEQ-500) run: 12.3M spots, 1.2G bases, 507.6Mb downloads | Figure 2E | 0 dilution_1 | SRX7990927 | Plasmid |
| 12 | BGISEQ (BGISEQ-500) run: 15.8M spots, 1.6G bases, 794.3Mb downloads | Figure 2E | 500 dilution_2 | SRX7990928 | Plasmid |
| 11 | BGISEQ (BGISEQ-500) run: 16.3M spots, 1.6G bases, 818.2Mb downloads | Figure 2E | 100 dilution_2 | SRX7990929 | Plasmid |
| 29 | BGISEQ (BGISEQ-500) run: 23.8M spots, 2.4G bases, 1.2Gb downloads | Figure 2E | 20 dilution_2 | SRX7990911 | Plasmid |
| 28 | BGISEQ (BGISEQ-500) run: 6.9M spots, 693.4M bases, 343.4Mb downloads | Figure 2E | 4 dilution_2 | SRX7990912 | Plasmid |
| 27 | BGISEQ (BGISEQ-500) run: 3M spots, 300.2M bases, 146.1Mb downloads | Figure 2E | 0.8 dilution_2 | SRX7990913 | Plasmid |
| 26 | BGISEQ (BGISEQ-500) run: 4.6M spots, 455M bases, 225.3Mb downloads | Figure 2E | 0.16 dilution_2 | SRX7990914 | Plasmid |
| 25 | BGISEQ (BGISEQ-500) run: 2.1M spots, 211.3M bases, 98.1Mb downloads | Figure 2E | 0 dilution_2 | SRX7990915 | Plasmid |
| 24 | BGISEQ (BGISEQ-500) run: 28.7M spots, 2.9G bases, 1.4Gb downloads | Figure 2E | 500 dilution_3 | SRX7990916 | Plasmid |
| 23 | BGISEQ (BGISEQ-500) run: 19.3M spots, 1.9G bases, 948.2Mb downloads | Figure 2E | 100 dilution_3 | SRX7990917 | Plasmid |
| 22 | BGISEQ (BGISEQ-500) run: 39M spots, 3.9G bases, 1.8Gb downloads | Figure 2E | 20 dilution_3 | SRX7990918 | Plasmid |
| 21 | BGISEQ (BGISEQ-500) run: 42.5M spots, 4.3G bases, 2Gb downloads | Figure 2E | 4 dilution_3 | SRX7990919 | Plasmid |
| 20 | BGISEQ (BGISEQ-500) run: 46.8M spots, 4.7G bases, 2.1Gb downloads | Figure 2E | 0.8 dilution_3 | SRX7990920 | Plasmid |
| 19 | BGISEQ (BGISEQ-500) run: 26.2M spots, 2.6G bases, 1Gb downloads | Figure 2E | 0.16 dilution_3 | SRX7990921 | Plasmid |
| 18 | BGISEQ (BGISEQ-500) run: 35M spots, 3.5G bases, 1.3Gb downloads | Figure 2E | 0 dilution_3 | SRX7990922 | Plasmid |
| 32 | ILLUMINA (Illumina MiSeq) run: 555,303 spots, 164.6M bases, 83Mb downloads | Figure 3 | CHLA2 | SRX8264270 | nasal swab |
| 33 | ILLUMINA (Illumina MiSeq) run: 1.3M spots, 384.9M bases, 201.7Mb downloads | Figure 3 | CHLA5 | SRX8264269 | nasal swab |
| 34 | ILLUMINA (Illumina MiSeq) run: 775,686 spots, 226.8M bases, 114.3Mb downloads | Figure 3 | CHLA3 | SRX8264268 | nasal swab |
| 35 | ILLUMINA (Illumina MiSeq) run: 1.2M spots, 364.9M bases, 188.1Mb downloads | Figure 3 | CHLA7 | SRX8264267 | nasal swab |
| 36 | ILLUMINA (Illumina MiSeq) run: 1.8M spots, 525.7M bases, 266.9Mb downloads | Figure 3 | CHLA11 | SRX8264266 | nasal swab |
| 37 | ILLUMINA (Illumina MiSeq) run: 870,090 spots, 256.9M bases, 130.8Mb downloads | Figure 3 | CHLA8 | SRX8264265 | nasal swab |
| 38 | ILLUMINA (Illumina MiSeq) run: 1.1M spots, 340.2M bases, 176.6Mb downloads | Figure 3 | CHLA13 | SRX8264264 | nasal swab |
| 39 | ILLUMINA (Illumina MiSeq) run: 332,823 spots, 100.5M bases, 51.2Mb downloads | Figure 3 | CHLA12 | SRX8264263 | nasal swab |
| 40 | ILLUMINA (Illumina MiSeq) run: 1.1M spots, 344.6M bases, 173.4Mb downloads | Figure 3 | MN908947.3 (Twist SARS-CoV-2 RNA) | SRX8264262 | not applicable |
| 41 | ILLUMINA (Illumina MiSeq) run: 710,098 spots, 214.4M bases, 110.6Mb downloads | Figure 3 | CHLA10 | SRX8264261 | nasal swab |
| 42 | ILLUMINA (Illumina MiSeq) run: 401,508 spots, 121.3M bases, 63Mb downloads | Figure 3 | CHLA6 | SRX8264260 | nasal swab |
| 43 | ILLUMINA (Illumina MiSeq) run: 945,198 spots, 285.4M bases, 144.1Mb downloads | Figure 3 | CHLA1 | SRX8264259 | nasal swab |
| 44 | ILLUMINA (Illumina MiSeq) run: 422,863 spots, 127.7M bases, 70Mb downloads | Figure 3 | CHLA4 | SRX8264258 | nasal swab |
| 45 | ILLUMINA (Illumina MiSeq) run: 1.3M spots, 401.2M bases, 203.1Mb downloads | Figure 3 | CHLA9 | SRX8264257 | nasal swab |

## Notes

### Competing Interest Statement

The authors have declared no competing interest.

